# Photizo: an open-source library for cross-sample analysis of FTIR spectroscopy data

**DOI:** 10.1101/2022.02.25.481930

**Authors:** Melissa Grant-Peters, Charlotte Rich-Griffin, Jonathan E. Grant-Peters, Gianfelice Cinque, Calliope A. Dendrou

## Abstract

With continually improved instrumentation, Fourier transform infrared (FTIR) microspectroscopy can now be used to capture thousands of high-resolution spectra for chemical characterisation of a sample. The spatially resolved nature of this method lends itself well to histological characterisation of complex biological specimens. However, commercial software currently available can make joint analysis of multiple samples challenging and, for large datasets, computationally infeasible. In order to overcome these limitations, we have developed Photizo - an open-source Python library for spectral analysis which includes functions for pre-processing, visualisation and downstream analysis, including principal component analysis, clustering, macromolecular quantification and biochemical mapping. This library can be used for analysis of spectroscopy data without a spatial component, as well as spatially-resolved data, such as data obtained via infrared (IR) microspectroscopy in scanning mode and IR imaging by focal plane array (FPA) detector.

## 1.0 Introduction

Fourier transform infrared microspectroscopy (µFTIR) enables non-destructive and label-free mapping of complex chemical information. The spatially resolved nature of this method lends itself well to the analysis of architecturally complex samples such as those of biological nature^[1]^. The functional groups specificity of µFTIR provide insight into biological queries, capturing biologically relevant molecules such as lipids, proteins, nucleic acids and carbohydrates^[2]^.

Continually improving instrumentation and spectral analysis methods are increasing the applicability of vibrational spectroscopy methods for disease characterisation and diagnosis. In particular, the chemical complexity captured in spectral features synergises well with machine learning approaches. These methods have repeatedly been shown to partition data based on these spectral features, distinguishing biochemical profiles of healthy control samples from pathological specimens^[3-5]^. In the context of histological characterisation with µFTIR specifically, clustering has been leveraged to distinguish the biochemical profile of different degrees of pathology within a given sample, with performance being comparable to a trained pathologist^[6]^.

However, performing this type of analysis while preserving spatial configuration data across multiple samples is challenging with currently available software. Commercially available software options, whilst rich in analysis functionality, can be computationally costly to run, often limiting processing to one sample at a time. When samples are processed and analysed individually with long running times for each step, this can increase the likelihood of errors and can be less systematic, thereby potentially compromising reproducibility. Quasar^[7]^, a recently available open-source spectroscopic data analysis toolbox extending the Orange suite, has overcome some of these challenges. However, its interactive interface comes at the cost of the capacity for cluster computing, which would be necessary for the analysis of large multi-sample datasets in a timely fashion.

### 1.1 Untapped potential: µFTIR enters the age of spatial high-resolution profiling technologies

Molecular profiling technologies which preserve spatial configuration data are becoming increasingly available, such as spatial transcriptomics, spatial proteomics and hyperplexed imaging methods. With multi-modal analysis approaches gaining prominence in the life and medical sciences to aid biological discovery and provide insights for patient prognosis^[8-10]^, diagnosis and therapy, an open-source tool for streamlined analysis of µFTIR data could unlock the significant potential of this method to better characterise the relatively understudied biochemical profile of tissues, and the data generated could then be integrated with other data modalities. This would substantially increase the utility of µFTIR data beyond data partitioning, and would also enable its use in disease characterisation since the multi-modal approach enables streamlined data exploration and validation. Integrative approaches would provide an avenue into inquiries of specific cellular and molecular processes linked to key macromolecular features.

In order to address this need, we present Photizo – an open-source Python library which makes use of functionality in the SCANPY library^[11]^ and AnnData objects to enable spectral analysis while preserving spectra-level clinical data annotation. It includes pre-processing, analysis and visualisation functions, including spatial mapping of spectra.

**Figure 1:**
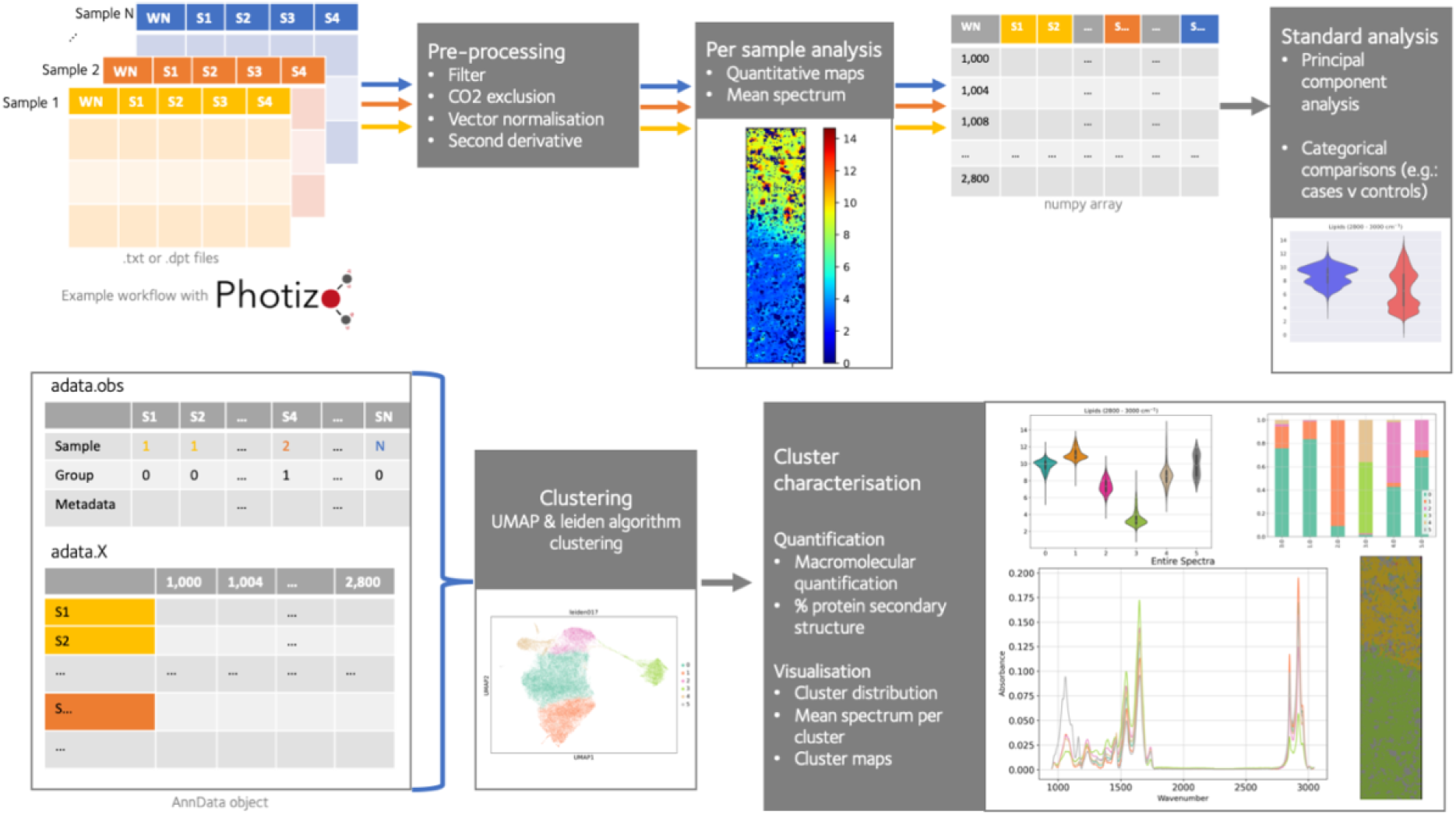
Example workflow of µFTIR data in Photizo.

## 2.0 Materials and methods

### 2.1 Inputs and pre-processing

Photizo takes as input .txt and .dpt files, where the first column contains wavenumbers and each subsequent column contains data points for a spectrum. This is read into a numpy array for pre-processing steps. Following pre-processing of each sample, subsequent steps can be performed for individual samples or for joint analysis of multiple samples. If using a data frame with multiple samples, we recommend creating an annotation data frame in pandas containing sample information (sample name, clinical data, etc). This will be necessary for visualisation of clinical variables as well as for partitioning into single-sample data for visualisation.

#### 2.1.1 Pre-processing

Photizo pre-processing includes resources for excluding outlier spectra with evidence of light scattering and spectra in regions with signal indicative of no sample (e.g.: sample holes, regions outside of sample borders), enabling application of vector normalisation to only include spectra of interest. In the process of hole exclusion, Photizo saves the positions of excluded spectra so that these positions can be repopulated prior to spatial mapping. We recommend spatially verifying the position of excluded spectra to ensure these are consistent with histological features (e.g. holes).

Pre-processing also enables the exclusion of the CO_2_ region – this is particularly useful when the CO_2_ captured is of atmospheric origin and does not contribute to the analysis in question. By excluding this region prior to clustering the user ensures that atmospheric alterations throughout the experiment do not contribute to clustering, creating batch effects. Calculating the second derivative of the spectra is also included in Photizo, which controls for baseline variation at the time of collection, thereby minimising batch effects in subsequent clustering.

### 2.2 Analysis

#### 2.2.1 Principal component analysis

Principal component analysis (PCA) can be used as a dimensionality reduction method prior to subsequent analysis and can be useful for identification of batch effects or spectral baseline effects prior to further analyses, as well as for discovering variables of genuine interest. Photizo has a PCA function optimised for spectral data, which rapidly generates visualisation of cumulative explained variance plots as well as a plot of the top 3 eigen-spectra. The PCA outputs can be used for other subsequent analyses, such as custom plotting and principal component projection.

#### 2.2.2 Clustering

Photizo includes clustering tools which make use of uniform manifold approximation and projection (UMAP) dimensionality reduction^[12]^ paired with the leiden algorithm^[13]^ for community detection, with flexibility regarding leiden algorithm resolution. Clustering may be performed with the entire spectra or only with a particular region of interest by using the functions to select regions of interest.

#### 2.2.3 Visualisation and quantification

Cluster profiling benefits from functions for visual spectral inspection which may include the entire spectra or exclusively particular regions of interest. Moreover, tools for quantitative comparisons also contribute to cluster characterisation, with functions implemented for numerical integration of the area below the spectra within the wavenumber window of interest. Selection of the window of interest may be verified with a specific spectral inspection function, where the user can account for subtle peak shifts in the data to select integration windows consistent with the collected data. Resulting quantified values can be compared across clusters using violin plot visualisation and be used for subsequent statistical comparisons.

Among the quantitative measures generated as outputs are estimates for secondary structure composition derived from the spectral features; these do not rely on spectral decomposition, but rather use statistically estimated content previously reported in the literature^[14]^, making this approach robust and reproducible.

Two key visualisation functions in Photizo enable spatial mapping of data in the configuration of data collection, requiring only the number of spectra collected in the x and y axes at time of collection. The first function maps integrated values for visualisation of chemical content estimation across the tissue for a particular region of interest. The second enables spatial mapping of cluster classification, which can later be compared to other characterisation of the tissue (e.g. histological staining).

### 2.3 Example workflow and reference dataset

In order to facilitate the use of the library by new users, we have made available infrared imaging by FPA detector data, with spatially resolved spectra collected from brain sections for exploration of the library’s functionality. This includes areas from three cases and three controls, thus enabling the user to perform a full workflow with reference figures, data and metadata prior to using the library on their own collected data.

## 3.0 Conclusions

Here we present Photizo, an open-source library for analysis of FTIR spectroscopy data, which includes functionality for analysing spatially resolved µFTIR data. This library is built in Python - a popular programming language with emphasised code readability - enabling users to analyse FTIR data with more flexibility regarding sample number and data size than currently available options, all at a low monetary cost. Photizo streamlines analysis of multiple samples, including the option of joint sample analysis, making its methods reproducible and easy to standardize across samples and datasets. Being built on Python, it can also be used for scripts submitted to cluster computing, vastly reducing computational costs for analysis. It has flexible functionality, facilitating reusability of basic functions and can be easily integrated into further workflows or analyses (e.g. statistical comparison of quantitative findings) as required by the user, and may also be adapted to the analysis of other vibrational spectroscopy methods, both spatially and non-spatially resolved (e.g.: Raman microspectroscopy, FTIR spectroscopy, etc). Importantly, while certain methodologies or tools utilised for Photizo come from biomedical sciences, the library is specimen-agnostic and can easily be used for spectral analysis of other sample types.

With the rise of integrative multi-modal analysis, this package contributes to closing the gap for µFTIR data to be analysed as part of larger integrative studies, providing biochemical context for other omics technologies. Jointly, these features contribute to maximising the utility of spectroscopy data with shorter processing times, at lower costs and with more flexibility than previous alternatives.

## Acknowledgements

This work was performed with support from the Wellcome Trust and the Interdisciplinary Bioscience DTP, supported by the BBSRC. Tissue samples and associated clinical and neuropathological data were supplied by the Multiple Sclerosis Society Tissue Bank, funded by the Multiple Sclerosis Society of Great Britain and Northern Ireland, registered charity 207495.

## Funding

This work was performed with support from the Wellcome Trust and Royal Society (204290/Z/16/Z) and the Interdisciplinary Bioscience DTP, supported by the BBSRC.

The code underlying this article, tutorials and instructions of how to access example data are available at https://github.com/DendrouLab/Photizo.

Conflict of interest: none declared.

## References

1. Baker, M.J., et al., Using Fourier transform IR spectroscopy to analyze biological materials. Nat Protoc, 2014. 9(8): p. 1771–91.

2. Bellisola, G. and C. Sorio, Infrared spectroscopy and microscopy in cancer research and diagnosis. Am J Cancer Res, 2012. 2(1): p. 1–21.

3. Heraud, P., et al., Early detection of the chemical changes occurring during the induction and prevention of autoimmune-mediated demyelination detected by FT-IR imaging. Neuroimage, 2010. 49(2): p. 1180–9.

4. Martel, C., et al., Diagnosis of idiopathic amyotrophic lateral sclerosis using Fourier-transform infrared spectroscopic analysis of patient-derived skin. Analyst, 2020. 145(10): p. 3678–3685.

5. Kneipp, J., et al., Detection of pathological molecular alterations in scrapie-infected hamster brain by Fourier transform infrared (FT-IR) spectroscopy. Biochimica et Biophysica Acta (BBA) -Molecular Basis of Disease, 2000. 1501(2): p. 189–199.

6. Wehbe, K., et al., Discrimination between two different grades of human glioma based on blood vessel infrared spectral imaging. Analytical and bioanalytical chemistry, 2015. 407(24): p. 7295–7305.

7. Toplak, M., et al., Quasar: Easy Machine Learning for Biospectroscopy. Cells, 2021. 10(9).

8. Eddy, S., L.H. Mariani, and M. Kretzler, Integrated multi-omics approaches to improve classification of chronic kidney disease. Nature Reviews Nephrology, 2020. 16(11): p. 657–668.

9. Miao, Z., et al., Multi-omics integration in the age of million single-cell data. Nature Reviews Nephrology, 2021. 17(11): p. 710–724.

10. Palla, G., et al., Spatial components of molecular tissue biology. Nature Biotechnology, 2022.

11. Wolf, F.A., P. Angerer, and F.J. Theis, SCANPY: large-scale single-cell gene expression data analysis. Genome Biology, 2018. 19(1): p. 15.

12. Becht, E., et al., Dimensionality reduction for visualizing single-cell data using UMAP. Nature Biotechnology, 2019. 37(1): p. 38–44.

13. Traag, V.A., L. Waltman, and N.J. van Eck, From Louvain to Leiden: guaranteeing well-connected communities.

14. Goormaghtigh, E., et al., Protein secondary structure content in solution, films and tissues: redundancy and complementarity of the information content in circular dichroism, transmission and ATR FTIR spectra. Biochim Biophys Acta, 2009. 1794(9): p. 1332–43.

